# Mammalian Uteri Contain a Diverse Virome: Insights from Healthy Mares

**DOI:** 10.1101/2025.02.26.640264

**Authors:** Lulu Guo, G. Reed Holyoak, Udaya DeSilva

## Abstract

The Earth’s estimated 10^31^ virions, primarily phages, significantly impact microbial ecosystems. Despite their abundance, viromes remain relatively understudied. This is especially the case in domestic animals. Although recent studies have described the presence of a dynamic commensal microbiome in mammalian uteri, no studies have attempted to characterize the commensal virome in a mammalian uterus. In this study we have established for the first time the presence of a sparse, but diverse native virome in the equine uterus. The resulting virome database consists of 513 non-redundant viral genomes (>2 kb). Taxonomic annotations revealed a diverse virome dominated by Gammaretrovirus, Mamastrovirus, Sapovirus and Rosenblumvirus. Notably, 75% of the assembled genomes represented novel species. The phylogenetic tree unveiled distinct clades suggesting unexplored viral diversity with the uterine environment. Furthermore, bacterial hosts for equine uterine phages were predicted, aligning with previous studies’ findings. Most notably, the study identified antibiotic resistance genes within the virome, hinting at potential gene transfer mechanisms between bacteria and viruses. This study establishes the first uterine virome of any mammal, shedding light on a previously unexplored domain. The findings highlight the potential for phage therapy in reproductive infectious diseases and the importance of understanding the maternal gestational environment. Moreover, the study underscores the need for further studies to expand the uterine viromes, paving the way to deeper understanding of uterine microbiome and its implications for animal and human health.

## Introduction

There are an estimated 10^31^ virions on earth^1^. Most of these viruses are thought to be phages that infect prokaryotes and compete with their hosts by at least an order of magnitude^2^. Based on the half-life of free viruses ∼(48h), it is estimated that 10^27^ viruses are produced per minute, and 10^25^ microorganisms - about 100 million metric tons – of bacteria are destroyed per minute^1^. Although bacterial and fungal biomes of most environments have been thoroughly investigated, viromes have not been deeply studied. Most sequence data in typical virome studies remain unidentifiable, and referred to as unexplored viral “dark matter”^3^. According to the National Center for Biotechnology Information (NCBI) Genomes database, only 11,673 complete viral genome sequences have been reported as of January 2024. This number represents a very small fraction of the total viral genetic diversity. Reproductive tract virome studies have focused mainly within the vagina because of accessibility and ease of sampling without cross contamination^4^. Additionally, the uterine environment has long been considered a sterile environment as early as Henry Tissier’s 1900 study^5^. However, this assumption started to be challenged in the mid-to-late 1980s, with reports indicating culturable bacteria from the uterus of healthy asymptomatic women^6789^. With the advent of high-throughput bacterial genomic analyses, many reports of uterine microbiomes of different species have been published, such as human^10^, dairy cow^11^, canine^12^, and equine^13^. Published virome studies on female reproductive tract mostly focuses on human vaginal environment, such as comparing the vaginal viromes of pregnant women with or without vaginitis^14^and the differences between health and dysbiotic vaginal DNA virome^15^ etc. Human vaginal virome analyses led scientists to believe that vaginal viromes are able to balance female genital tract bacteriome, mucosal immunity, sexual and reproductive health and disease^16^, and that change of cervicovaginal DNA virome will affect genital inflammation and microbiota composition^17^. However, understanding the role of virome in animal gynecological health is also important because they are an essential part of the entire ecosystem. Our previous metagenomic analyses of the equine uterine microbiome suggested the existence of a commensal virome. Here we report the identification of a native virome of the equine uterus, the host prediction for those viruses, and the presence of antibiotic resistance in the uterine environment. To the best of our knowledge, this is the first comprehensive analysis of the uterine virome of a mammal.

### Equine Uterine Virome Database

We obtained 272,710,338 clean reads (∼4 billion nucleotides with an average read length of 140 bp) with Phred scores higher than 40. These assembled into 32,084 contigs with an average lengt∼h 550 bp. Minimum contig length wa∼s 200 bp. 2789 contigs were over 10 kb in length. 1750 of the larger contigs were determined to be high-quality for containing majority (>70%) of the organism’s predicted genome^18^. The N50 length, defined as the shortest contig length that covers at least half of the total base content was ∼500 bp and the N90 length defined as the shortest contig length that ensures that the contigs of that length, or longer contains at least 90% of the sum lengths of the contigs was∼310 kb (data not shown). The GC content of the contigs were ∼40% GC and none of them had gaps. 66 contigs were complete circular genomes. No detectable DNA was observed in either negative control in a bioanalyzer and were not sequenced. To predict viral sequences, all the contigs were filtered through DeepVirFinder^19^, which takes advantages of deep learning to build convolutional neural networks to identify viral contigs without using reference databases 462 contigs were recognized as virus. VirSorter^20^, which relies on comparing sequence to known virus and viral-like features was also used to identify viral contigs and 78 contigs were identified. Twenty-seven of these contigs were recognized by both tools. We then filtered sequences at a 95% average nucleotide identity (ANI) threshold and 70% aligned fraction threshold obtaining a database of 513 uterine viral sequences, referred to here as the equine uterine virome database.

### Equine Uterus Harbors Diverse Range of Novel Virome Sequences

Taxonomic annotations of the contigs were performed by CAT^21^, which classifies contigs according to genes, open reading frames (ORFs) prediction, and protein mapping. The IMG/VR database was used during annotation. The most plausible abundance in different levels based on viral contigs is shown in Figures 2(a)-2(f). The mare endometrium is dominated by Gammaretrovirus, Mamastrovirus, Sapovirus and Rosenblumvirus at genus level. Gammaretrovirus, Mamastrovirus and Sapovirus are RNA viruses while Rosenblumvirus is a dsDNA virus. Most abundant predicted Phyla are depicted in Figure 3. As shown, 11 classes, 11 orders, 14 families, 8 genera, 7 phyla and 29 species were identified. To represent richness, evenness, and statistical significance (p-adj <0.05), the alpha diversity was calculated by R vegan package. Alpha rarefaction of observed metrices showing species richness in samples reached near-saturation, but it only considers observed OTUs. However, chao1 considers not only observed OTUs, but also non-observed rare species. The results show the sequence coverage is sufficient. The non-parametric Kruskal-Wallis test was used for Shannon index and Simpson index (Figures 4(a) & 4(b)). Simpson index focuses more on relative abundance while Shannon index focuses more on species richness. Alpha diversity indices at different phylogenetic classifications are shown in Table 1.

**Table 1.**
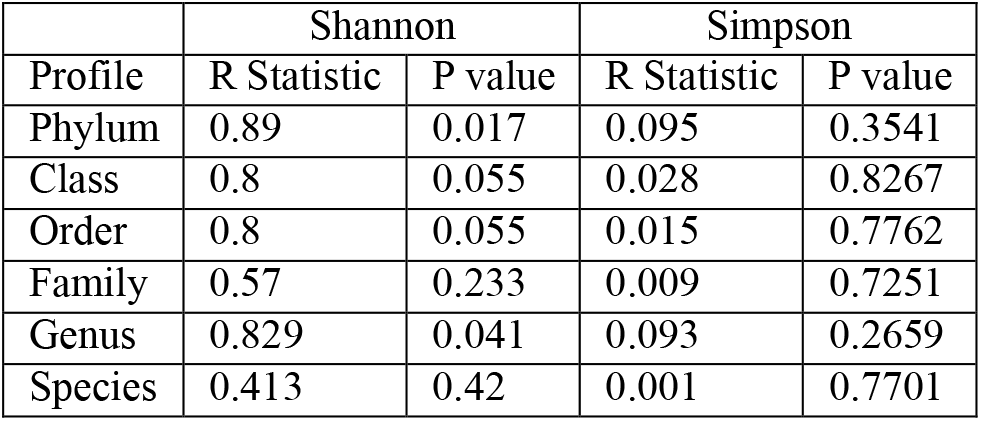
Alpha diversity indices at different phylogenetic classifications.

| Profile | Shannon |  | Simpson |  |
| --- | --- | --- | --- | --- |
|  | R Statistic | P value | R Statistic | P value |
| Phylum | 0.89 | 0.017 | 0.095 | 0.3541 |
| Class | 0.8 | 0.055 | 0.028 | 0.8267 |
| Order | 0.8 | 0.055 | 0.015 | 0.7762 |
| Family | 0.57 | 0.233 | 0.009 | 0.7251 |
| Genus | 0.829 | 0.041 | 0.093 | 0.2659 |
| Species | 0.413 | 0.42 | 0.001 | 0.7701 |

**Figure 1.**
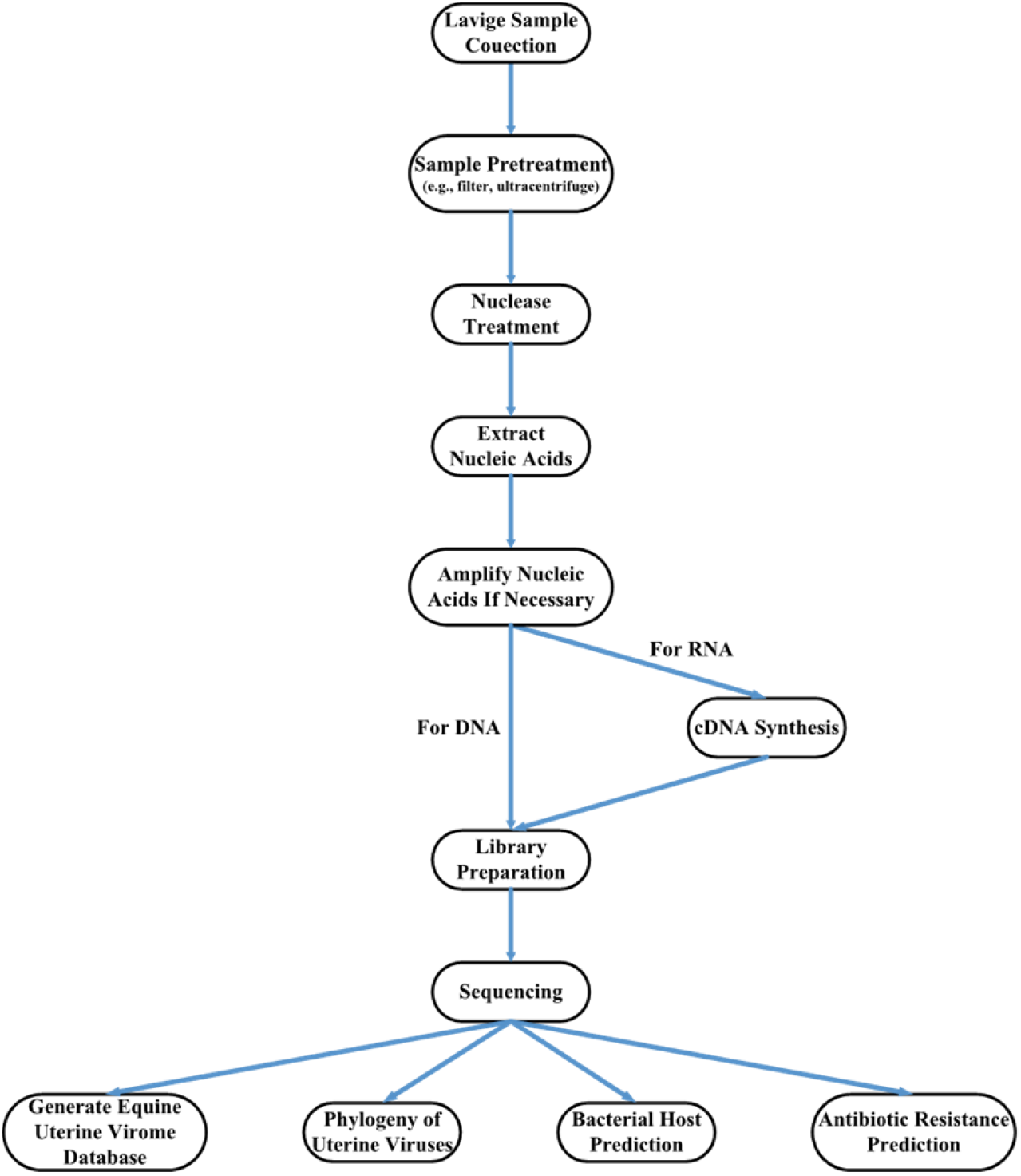
Overview of experimental workflow.

**Figure 2.**
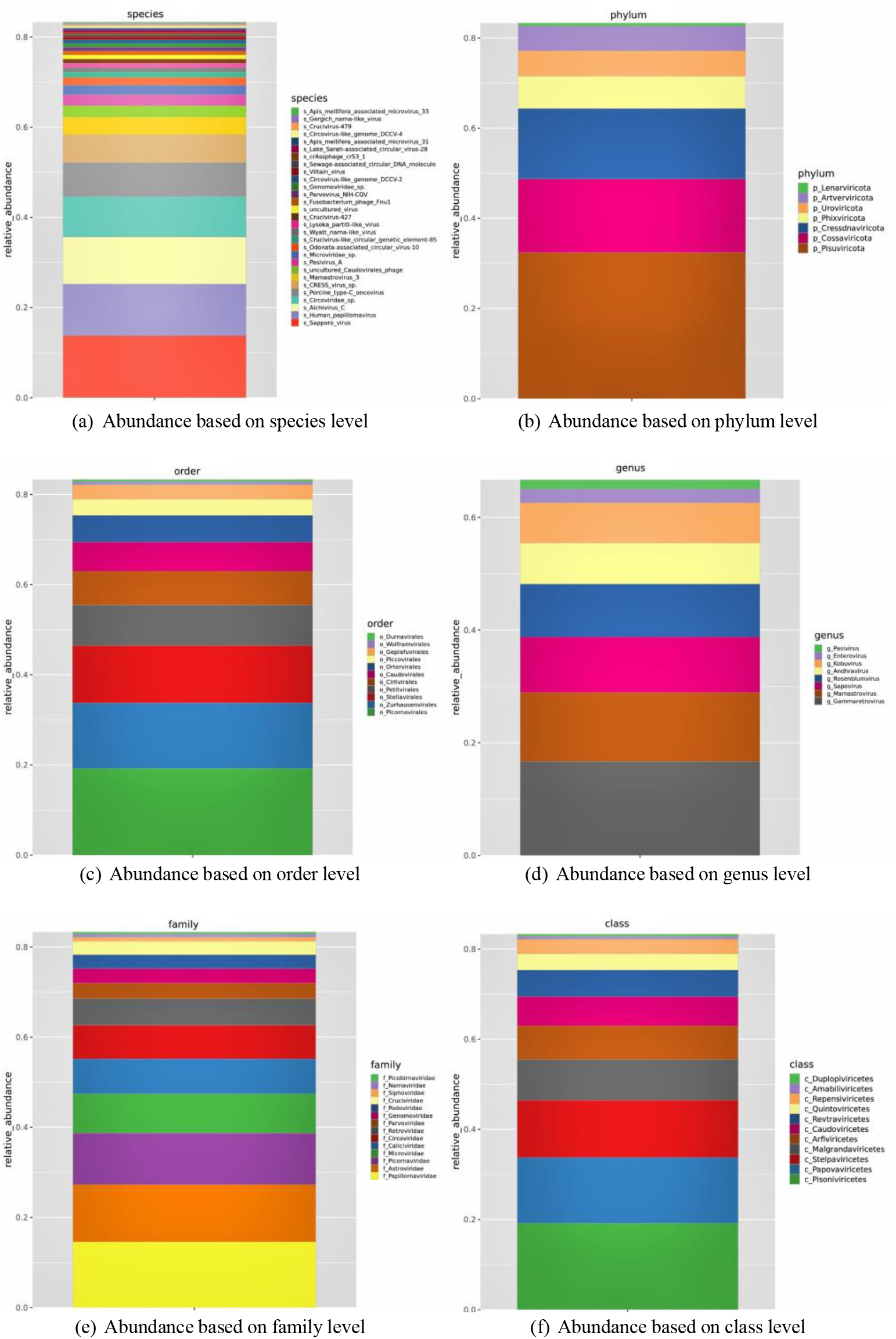
The abundance profiles are depicted across various taxonomic levels, ranging from phylum to species. Each figure provides insight into the distribution and prevalence of taxa within the analyzed dataset, shedding light on the taxonomic composition and diversity present. Through comprehensive examination at different taxonomic resolutions, from broad classifications such as phylum down to specific species, these figures offer a nuanced understanding of the microbial community structure within the uterus. The dominance of Gammaretrovirus, Mamastrovirus, Sapovirus, and Rosenblumvirus at the genus level in the mare endometrium underscores the diversity of viral populations within this biological niche. Notably, Gammaretrovirus, Mamastrovirus, and Sapovirus are RNA viruses, whereas Rosenblumvirus belongs to the dsDNA virus group.

**Figure 3.**
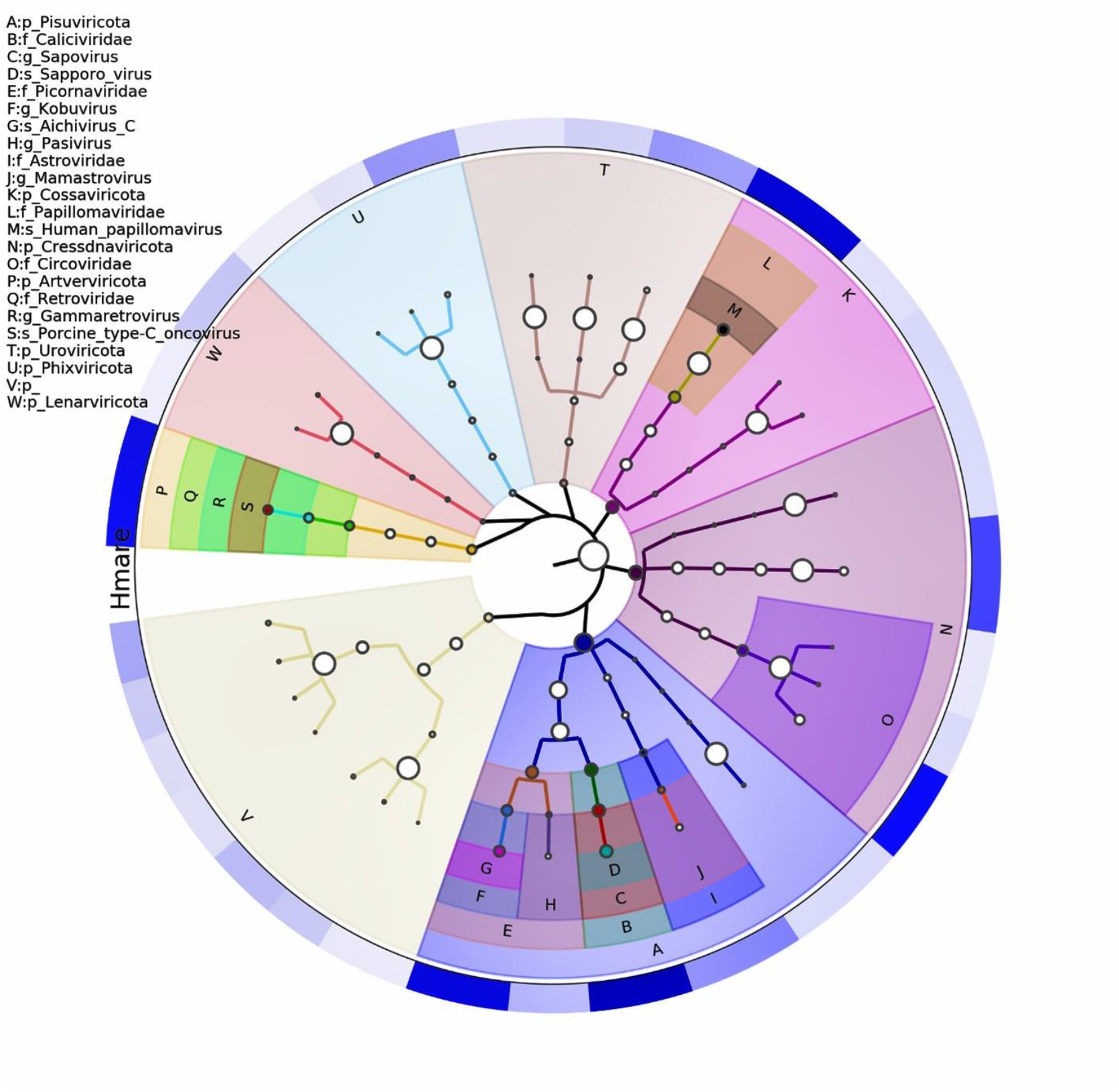
A contig-based abundance analysis unveils a detailed representation of the genomic content within the assembled sequences. Additionally, the identification of 11 classes, 11 orders, 14 families, 8 genera, 7 phyla, and 29 species underscores the taxonomic richness and diversity present in the microbial community.

**Figure 4.**
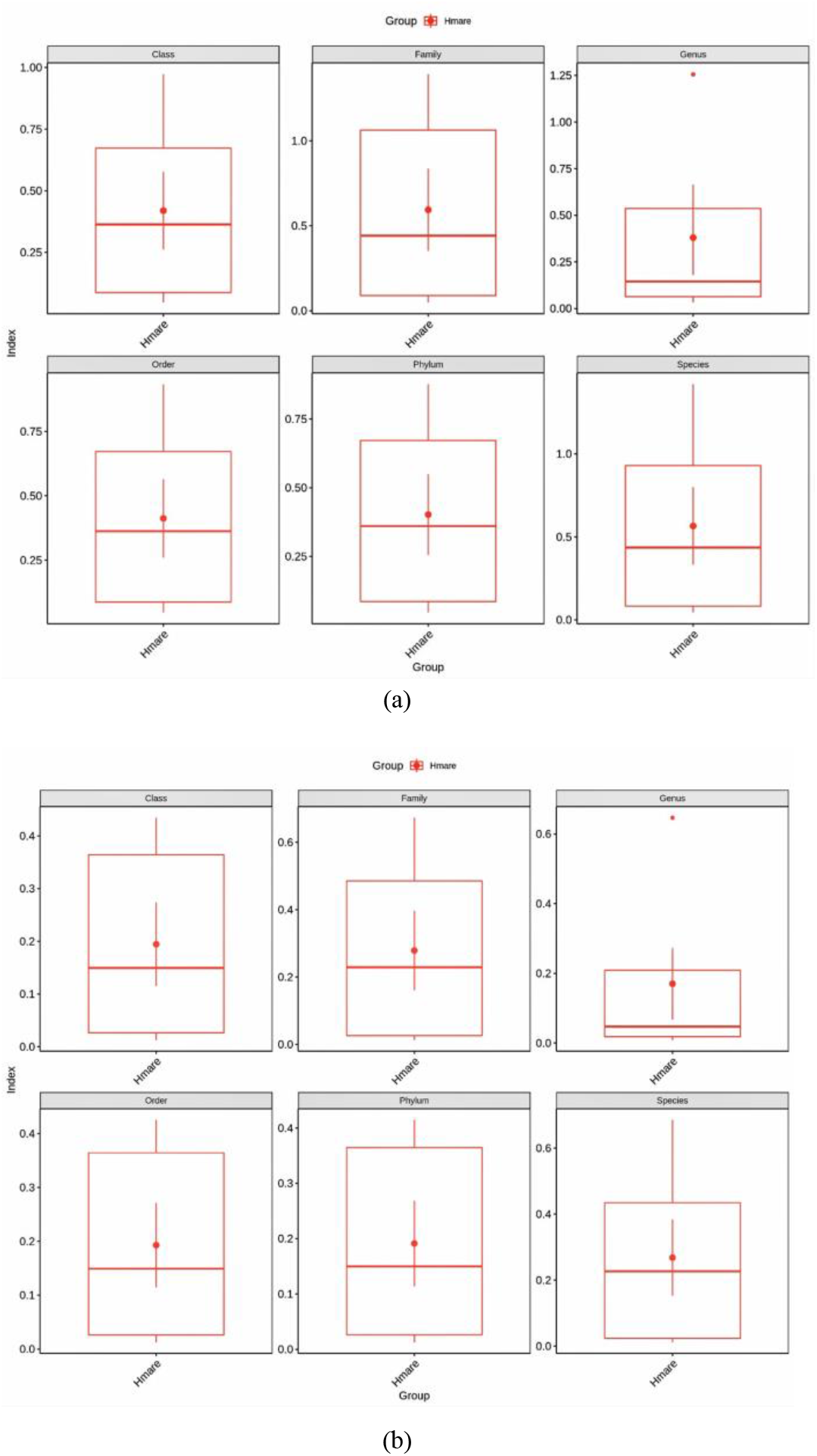
(a) The alpha diversity of the analyzed ecosystem was assessed using the Shannon index. This metric provides a comprehensive measure of biodiversity within a specific community, taking into account both the abundance and evenness of different taxa. A higher Shannon index value signifies greater diversity, reflecting a more balanced distribution of species and indicating a richer and more complex ecological landscape. (b) The alpha diversity of the studied ecosystem is evaluated through the Simpson index. This index offers a complementary perspective on biodiversity, focusing on the dominance or concentration of species within the community. A lower Simpson index indicates higher diversity, emphasizing the presence of a more evenly distributed array of species.

### Phylogeny of Uterine Viruses

To discover more details about viruses within the uterine lumen, we performed comprehensive virome profiling of the uterine samples by mapping 2789 contigs against the NR-V database to make sure we covered prophages. 513 previously predicted viral contigs were utilized to build a phylogenetic tree by MATLAB software and ITOL^22^. Two main branches were generated, and trees were built for both (data not shown). This further indicates there are many unknown viral species in the uterine environment. The relationships between some viruses are close, suggesting that they probably evolved from the same organism or mutated into a new virus.

### Bacterial Host Prediction for Equine Uterine Phages

We predicted the most likely bacterial host for the phages we obtained from equine uteri. First, we screened our contigs with the CRISPRCasdb spacer database to select all potential host information. Then, the phage-bacteria interaction database, Microbe Versus Phage (MVP) was used to arrange the similarities between contigs and known virus sequences. Finally, correlation was applied to predict the bacterial host for the contigs. The core hosts we found were Clostridiales at order level, and Streptococcus and Staphylococcus at genus level, which parallels our previous equine uterine microbiome study^13^.

### Presence of Antibiotic Resistance Genes in Equine Uterine Virome

Investigating the uterine viral sequences further, we employed a method based on ORFs annotation for subsequent analysis of related community functions^23^. Using BWA-MEME^24^, the effective data of each sample after removing the host sequence were compared to the initial ORFs set of each sample. The number of reads of a gene in each sample were calculated. According to the comparison results, ORFs of each sample were filtered. Dereplication was performed with CD-HIT^25^ using global identity threshold of 99%. A total of 463 ORFs were predicted by Prokka^26^. Antibiotic resistance genes were annotated by Comprehensive Antibiotic Research Database (CARD) and Resistance Gene Identifier^27^. Although antibiotics do not have direct effect on viruses, studies have found that viromes from a variety of environments carry antibiotic resistance genes^28^. As suggested by multiple studies^29,30^, bacteriophages may play a role in horizontal transfer of genes conferring antibiotic resistance to bacteria. Therefore, it is particularly important to detect drug resistance factors within viral functional genes. The core antibiotic resistance genes detected in the uterine viruses were gene 88 and gene 163, being involved with resistance to one or more antibiotics (Figures 5(a) & 5(d)) by target modification mechanisms (Figures 5(b) & 5(e)). The most predicted antibiotic resistance is to lincosamides, macrolides, and streptogramin (Figure 5(c)).

**Figure 5.**
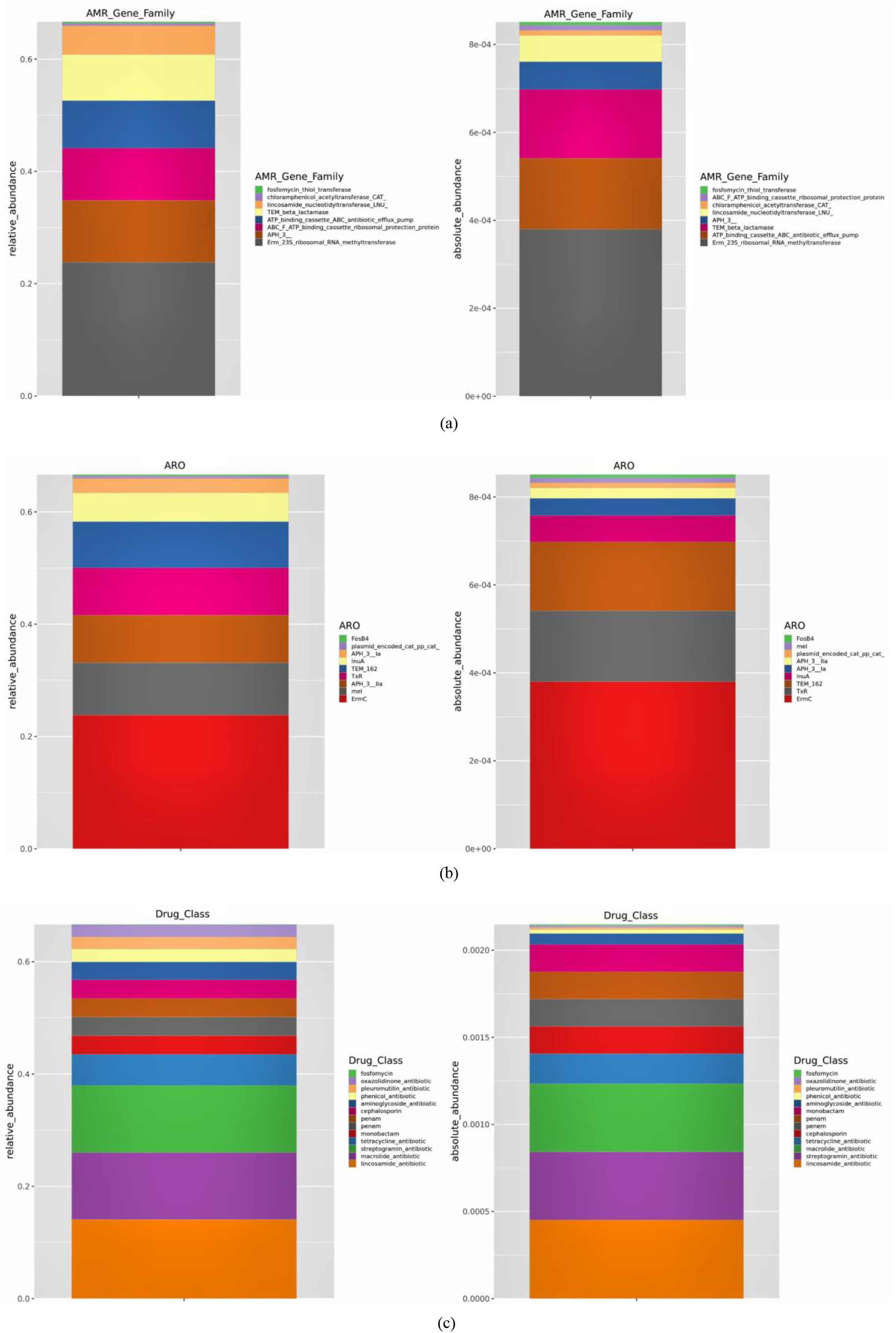
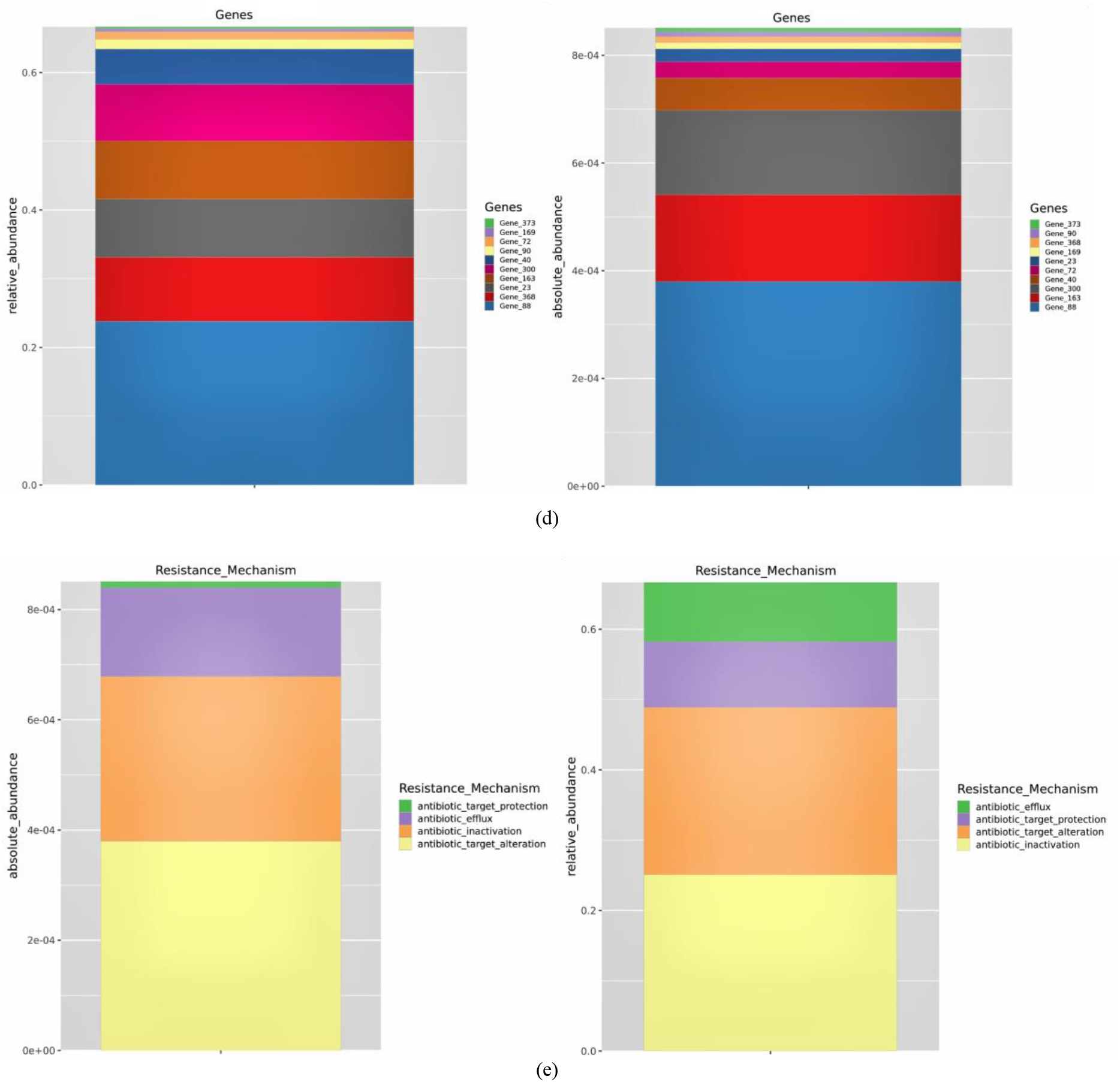
(a) The resistance mechanism of antibiotic genes illuminating the strategies employed by these genes to confer resistance to antimicrobial agents. The identification of core antibiotic genes in uterine viruses associated with target modification mechanisms indicates a prevalent strategy within the microbial community to evade the effects of antibiotics. (b) The abundance of antibiotic genes within the viral genome is highlighted, providing a comprehensive view of the prevalence and distribution of these resistance determinants among viral elements. The identification of core antibiotic genes, specifically gene 88 and gene 163, within the uterine viruses signifies their prominent role in conferring antibiotic resistance within the microbial community associated with the uterus. (c) An exploration is undertaken to identify the most likely resistant drugs associated with the antibiotic genes under consideration. The most predicted antibiotic resistance in the study is to lincosamides, macrolides, and streptogramin, highlighting the prevalence of resistance mechanisms against these classes of antibiotics within the analyzed microbial community. (d) Antibiotic Resistance Ontology. (e) An exploration is undertaken to identify the most likely resistant drugs associated with the antibiotic genes under consideration. The most predicted antibiotic resistance in the study is to lincosamides, macrolides, and streptogramin, highlighting the prevalence of resistance mechanisms against these classes of antibiotics within the analyzed microbial community.

## Discussion

To date, genome sequencing has made remarkable contributions to the fields of microbiology, biology, and physiology, leading some to state that all organisms are metaorganisms^31^. The microbiota, microbial communities within and on the animal body, are regarded as an integral part of the metaorganism blueprint and play important roles in determining the health of animals^4^. The population and diversity of the virome has a major impact on the community of hostile microbes and pathobionts. In this study, we have obtained and analyzed more than 500 high-quality equine uterine viral contigs. To the best of our knowledge, we present the first uterine virome in a mammal. As expected, majority of the viral contigs (n=462) were identified using DeepVirFinder^19^ that predicts viral sequences using deep learning and building convoluted neural networks without relying on reference databases. We identified 78 contigs using VirSorter, that relies on comparing sequences to known virus and virus-like sequences. Only 27 contigs were identified by both programs. As it is estimated that a vast majority of viruses that exist are yet to be identified^32^, this result was expected and was the experience of a few other studies^33,34^. The uterus has traditionally been considered a sterile environment. Although being far less robustly populated than the gut or the vagina, uterine microbiomes have been previously described. Our approach (Figure 1) allowed us to capture viral genomes from this comparatively sparse, yet more diversly colonized environment. Since the uterus is relatively understudied, we were unable to classify many contigs below Family level, although we have assigned several contigs to different genera and species, they represented only a fraction of the identified viruses. Over 75% of the assembled genomes represented novel species. The uterine endometrium hosts a highly diverse and novel viral population. The most abundant family found is Papillomavirus, a double strand DNA virus. This virus has previously been reported in cutaneous lesions on the horse^35^, and this is the first report of it being found in the uterus. It should be noted that this virus, and closely related (almost identical) bovine papilloma virus, have been associated with sarcoid, squamous cell carcinoma/genital papillomatosis in the horse^36^ and associated with similar lesions in the human uterine environment^37^. This is the first time the single strand RNA virus families of Astroviridae and Caliciviridae, have been identified in horses. We believe there are still many other viruses within the uterus remaining to be identified and studied. There are also abundant phage families that have been discovered in this study, such as Microviridae, Siphoviridae, Podoviridae and others, which suggest the possibility of the use of these native phages in targeted non-antibiotic therapy of reproductive infectious diseases in the future. This opens the door to further studies in this area. Multiple Displacement Amplification (MDA) that was used to amplify viral DNA is known to preferentially amplify certain DNA sequences over others^38,39^. Consequentially, the relative composition of different viral families identified in this study may have been influenced by our methodology. We limited our samples to healthy animals at a single stage of estrous cycle that minimizes ascending migration of vaginal microorganisms. We only sampled 12 animals and they were all from the same facility and in the same geographical area. As such, we are certain that the uterine virome of horses is much more diverse than is reported in this study. Our goal was to establish that a diverse and complex native virome exists in a mammalian uterus and we have successfully accomplished that goal. We were surprised by the prevalence and diversity of antibiotic resistance genes we found in viral genomes. Although having antibiotic resistance genes does not directly affect the virus, antibiotic disease gene prevalence among viral genomes has been observed before^28,29,30^. Although it was outside the scope of this work, checking if any of the uterine bacteria had acquired the same resistance elements would be interesting. Antibiotic resistant microbes are threatening the treatment of infectious diseases. At present, the identification of drug resistance genes can help us understand drug resistance mechanisms and provide references for disease treatment and drug development. Our study reveals that antibiotic resistance genes also reside in viral genomes. The predicted resistance mechanism of the antibiotic gene family reflects the co-evolution between bacteria and viruses. This suggests the existence of gene flow networks between phylogenetically distinct bacterial and viral species in the uterus. In this study, we also found that the uterine viral contigs were separated into 2 main clades. This result highlights the importance of building a well-established uterine virome database, as the combined effect sizes of highly related phage genomes can help reveal the association of specific clades with their bacterial hosts and animal health^40^.

## Methods

Topical subheadings are allowed. Authors must ensure that their Methods section includes adequate experimental and characterization data necessary for others in the field to reproduce their work.

### Animals and Sample Collection

Twelve healthy mares from Oklahoma were selected for sample collection in this study. Mares were between 3 and 14 years of age, with clinically normal reproductive tracts. Mares were sampled in a single day during the normal breeding season (mid-spring in the northern hemisphere) during the luteal phase of their estrous cycle (diestrus) between day 7 and 10 post-ovulation. At this time, under the influence of progesterone the cervix is closed and sulfomucins significantly reduces the potential for transcervical migration of vaginal bacteria. All mares had the same diet for at least 2 months before sample collection. None of the mares had recent antiviral or antibiotic exposure. All mares had uterine luminal endometrial microbiota samples collected via 150 ml of saline infused through a triple guarded system, where the sample collection catheters were protected from the vestibular and vaginal environment and brought to the external os of the cervix inside a sterile disposable vaginal speculum. The sample collection catheter was a stallion urinary catheter (Jorgensen Labs, Loveland, CO, USA) inserted within a 36 Fr. 70 cm equine embryo flush catheter (IMV Technologies, Holland, MI, USA) which served as the second guard to prevent contamination from the cervical external os and lumen. The embryo flush catheter was inserted into the cervix to the level of the internal cervical os, the balloon inflated with∼50 mL of air to seal the cervix, and then the urinary catheter advanced into the uterine lumen. Saline was infused, massaged within the lumen, and collected from the uterus via gravity flow. A mid-stream catch of approximately 50 mL was collected into sterile 50 mL conical tubes and immediately capped and placed on ice. The urinary catheter was then retracted into the flush catheter, and both retracted into the speculum, before removal back through the vaginal vault. This small volume lavage sampling technique has been shown to have the highest return rate for microbial collection^41^. We enhanced the guarded protection from potential vestibulovaginal contamination by adding the disposable speculum to the collection process. All samples were collected on a single day in the same facility and processed together to eliminate a batch effect. A saline sample was flushed through the catheter at the time of collection as well as a second aliquot of the same batch of saline was processed alongside regular samples throughout the experimental protocol as negative controls. Animal use and sample collection procedures were approved by the Oklahoma State University Institutional Animal Care and Use Committee. An overview of the experimental workflow described in this manuscript is provided in (Figure 1).

### Viral Nucleic Acid Isolation and Amplification

Lavage samples were decanted and sequentially filtered through a 0.45 *µ*m and 0.22 *µ*m filter by using Vacuum Driven Sterile Filter (Millipore Steriflip, Burlington, MA, USA). Viral particles were enriched by using ultracentrifuge and Pellet Paint^®^ Co-Precipitant (Novagen, Madison, WI). Viral enrichments were nuclease treated at 37°C for 90 minutes to remove contaminating free floating DNA and RNA. Nucleic acid was extracted from viral enrichments using the QIAamp UltraSens Virus Kit (Qiagen, Germantown, MD, USA) and following the manufacturer’s protocol. Quantifying the purified Viral nucleic acids was done using a NanoDrop ND-1000 spectrophotometer (NanoDrop Technologies, Wilmington, DE, USA). A 25-ng aliquot of purified nucleic acid was used as template for amplification using the REPLI-g Mini Kit (Qiagen, Germantown, MD, USA). Nucleic acid quantification was done using a Qubit 3.0 Fluorometer (Thermo Fisher Scientific, Waltham, MA, USA). Amplified samples were kept at 4°C for short-term storage.

### cDNA Synthesis and Library Preparation

Up to 1 *µ*g template RNA and ProtoScript^®^ II First Strand cDNA Synthesis Kit (New England Biolabs, Ipswich, MA, USA) were used in first strand cDNA synthesis following kit protocols. The second strand cDNA synthesis was done using NEBNext Ultra II Non-Directional RNA Second Strand Synthesis Module (New England Biolabs Inc.) according to the kit protocols. Metagenomic library preparation was performed using the NEBNext Ultra II Library Prep Kit (#E7645L) and Index Primers (#E7500S and #E7335L) from New England Biolabs. The protocol associated with these kits was used. Briefly, each library sample was quantified and run on a bioanalyzer to establish library molarity (nM) for sequencing. Samples were diluted to equivalent molarities and pooled for sequencing via Illumina HiSeq 2500 platform. The raw sequencing data used in this study is available at the Sequence Read Archive (SRA) of NCBI under the Project number PRJNA1122187 and accession numbers SAMN41777987-993.

### Quality Control and Metagenomic Assembly

To ensure data quality and avoid erroneous results, sequencing data needs to be quality filtered and preprocessed before use in subsequent analyses. FastQC^42^ software was used to evaluate the quality of sequencing data. Then, a common NGS quality control tool, Trimmomatic^43^ was used for pretreatment including filtering low-quality reads, removing low-quality bases and unrecognized base (N) at the end of reads, removing sequencing adapters, removing short sequences etc. In order to avoid host contamination, sequence were further quality-filtered with the latest equine database EquCab3 by using Bowtie 2^44^ and Samtools^45^ to remove the host sequences. MetaSPAdes^46^ was then used to assemble the paired-end reads.

### Viral Sequence Prediction

VIBRANT^18^ was used to analyze the contigs larger than 10kb and predict viral sequences since it can process large metagenomic assemblies based on combined machine learning and protein similarity approach. VirSorter^20^ was used to identify the virus contigs and to detect prophages by aligning a clean read with the NT (Nucleotide Sequence Database) library of non-redundant sequences through BLAST and aligned sequences that are homologous to viruses and bacteria. Groups included were dsDNAphage, NCLDV, RNA, ssDNA and lavidaviridae. A minimum score of 0.5 was used. Category 1, 2, 4, 5 were considered from VirSorter prediction. DeepVirFinder^19^] was used for deep learning to distinguish viral from bacterial contigs. Contigs with score > 0.9 and p-value < 0.01 were considered from DeepVirFinder prediction.

### Contigs Annotation, Clustering and Taxonomy

Viral genomes were annotated with Prokka^26^] by using viral kingdom. Based on identifying coordinates of genomic features within contigs, Prokka can annotate genes and save the information in standards-compliant output files. vConTACT2^47^ was used to form clusters by creating mapping file as it has been widely used to group phage sequences into clusters roughly corresponding to the genus level. Comparative Annotation Toolkit (CAT)^21^, using long read for taxonomic classification was used to provide taxonomic context of contigs. The function of the dereplicated genes were annotated by eggNOG database^48^, KEGG^49^ and CAZy database^50^. Kraken2 and Bracken was used to calculate species abundance^51^. Abundance was calculated by first classifying the sequence using Kraken2 and then estimating the abundance of each species with Bracken. Contig-based analyses were performed using Contig Annotation Tool (CAT)^52^.

### Bacterial Host Prediction and Antibiotic Resistance Gene Detection

Bacterial hosts were predicted by using the spacer sequence library from the CRISPRCasdb database and conducting a blastn-short comparison on assembled contig sequences to identify potential phage hosts^53^. Microbe versus Phage (MVP) database was used to build the database of phage-prokaryotic interactions^54^.

## Acknowledgements (not compulsory)

Acknowledgements should be brief, and should not include thanks to anonymous referees and editors, or effusive comments. Grant or contribution numbers may be acknowledged.

## Author contributions statement

Must include all authors, identified by initials, for example: A.A. conceived the experiment(s), A.A. and B.A. conducted the experiment(s), C.A. and D.A. analysed the results. All authors reviewed the manuscript.

## Additional information

To include, in this order: **Accession codes** (where applicable); **Competing interests** (mandatory statement).

The corresponding author is responsible for submitting a [competing interests statement](http://www.nature.com/srep/policies/index.html#competing) on behalf of all authors of the paper.

This statement must be included in the submitted article file.

